# A plug-and-play system for enzyme production at commercially viable levels in fed-batch cultures of *Escherichia coli* BL21 (DE3)

**DOI:** 10.1101/263582

**Authors:** Sujata Vijay Sohoni, Paras Harendra Kundalia, Adarsh G. Shetty, Avinash Vellore Sunder, Raghavendra P. Gaikaiwari, Pramod P. Wangikar

## Abstract

Commercial exploitation of enzymes in biotransformation necessitates a robust method for enzyme production that yields high enzyme titer. Nitrilases are a family of hydrolases that can transform nitriles to enantiopure carboxylic acids, which are important pharmaceutical intermediates. Here, we report a fed-batch method that uses a defined medium and involves growth under carbon limiting conditions using DO-stat feeding approach combined with an optimized post-induction strategy, yielding high cell densities and maximum levels of active and soluble enzyme. This strategy affords strict control of nutrient feeding and growth rates, and ensures sustained protein synthesis over a longer period. The method was optimized for highest titer of nitrilase reported so far (247 kU/l) using recombinant *E. coli* expressing the *Alcaligenes* sp. ECU0401 nitrilase. The fed-batch protocol presented here can also be employed as template to produce a wide variety of enzymes with minimal modification, as demonstrated for alcohol dehydrogenase and formate dehydrogenase.

## Introduction

Enzyme based biocatalytic processes have great potential in the production of pharmaceuticals especially involving chemical, regio- or stereo-selective reactions under mild reaction conditions (Buccholz et al. 2012). Nitrilases are hydrolases that can convert nitriles to enantiopure carboxylic acids, which have important applications as pharmaceutical intermediates or agrochemicals (Gong et al. 2012). Commercially viable processes have been established using nitrilases for the production of (R)-mandelic acid (Xue et al. 2013), key intermediate for the synthesis of anti-obsesity and anti-tumour agents, and (R)-o-chloromandelate (Wang et al. 2013), a key intermediate for synthesis of platelet aggregation inhibitor Clopidogrel, with high yields and optical purity. Alcohol dehydrogenases (ADH) represent another class of enzymes which have been exploited commercially, chiefly in the asymmetric reduction of ketones to optically active alcohols, which are intermediates in the synthesis of statins and anti-cancer drugs (Hall and Bommarius 2011).

The commercial viability of these processes is dependent on the availability of the biocatalyst; hence an economical method for the large-scale production of soluble and active enzyme in high yields is a major requirement. To date, *E. coli* based expression systems remain the most commonly used for production of microbial enzymes (Rosano and Ceccarelli 2014) due to the ease of cloning and genetic manipulation, and the availability of multiple engineered production strains feasible for fast cultivation. Routine lab scale expression often involves strains like BL21 (DE3) and pET plasmid vectors, which are based on a strong promoter (T7) and an operator (*lac*) (Terpe 2006). In the context of enzyme production in the industry for developing commercial biotransformation processes, it is prudent to grow the recombinant *E. coli* to high cell densities, while monitoring the rate of growth and protein synthesis to achieve maximum enzyme titres. This necessitates the development of a carefully optimized fermentation/bioprocess strategy.

While fed-batch fermentation is a prerequisite for cultivation of *E. coli* at high cell densities, selection of a proper feeding strategy is critical for inducible protein expression (Shiloach and Fass 2005). Feed-forward methods include feeding at a constant rate, and exponential feeding that allows the cells to grow at a controlled specific growth rate (μ) using the carbon source as a limiting nutrient (Korz et al. 1995). Other methods work based on feedback control of carbon source utilization reflected by changes in pH (pH-stat) or dissolved oxygen (DO-stat, Castan and Enfors 2000), respectively. The selected approach should be able to achieve strict control of the growth rate, and avoid overfeeding and accumulation of inhibitory products such as acetate or organic acids (Schaepe et al. 2014). In addition, other parameters including temperature and the time of induction also need to be optimized in tandem, as uncontrolled high level protein expression at high cell densities could burden the metabolic capabilities of the cell, leading to growth inhibition, protein misfolding or formation of inclusion bodies despite high protein yields (Choi et al. 2006).

The present report details the development of a cost-effective fed-batch method based on a DO-stat feeding approach, for achieving high cell densities and significantly high enzyme titers in cultivation of recombinant *E. coli* BL21(DE3) on fully defined mineral medium. Different process parameters including nutrient feeding, oxygen transfer rate and induction conditions were optimized for the growth of *E. coli* expressing nitrilase from *Alcaligenes* sp. ECU0401 (Zhang et al. 2010). Using the optimized strategy, we could achieve maximum nitrilase yields reported so far (247 kU/1). The process was also administered with minimal changes to achieve high enzyme titers of two other industrially relevant enzymes – ADH from *Lactobacillus kefir* and formate dehydrogenase (FDH) from *Candida boidinii*), suggesting its potential applicability for the production of a wide range of enzymes in the *E. coli* BL21/pET system.

## Experimental methods

### Chemicals

All the chemicals and solvents used in this study were procured from Merck (Germany) unless mentioned otherwise and were of analytical grade and HPLC grade, respectively. Carbenicillin and isopropyl β-D-1-thiogalactopyranoside (IPTG) were procured from Sigma Aldrich (USA).

### Bacterial strain, media and culture conditions

*E. coli* BL21 (DE3) strain (Novagen, Billerica, MA) was used as the expression host. Custom synthesized and codon optimized genes for (1) nitrilase from *Alcaligenes* sp. ECU0401 (UniProt ID C5IIS9), (2) ADH from *Lactobacillus kefir* (UniProt ID Q6WVP7) and (3) FDH from *Candida boidinii* (UniProt ID O13437) were procured from Biomatik Corporation (Ontario, Canada). All genetic manipulations were carried out in *E. coli* DH5α strain. Each of the enzyme genes was first sub-cloned in pET21a vector (Novagen, Billerica, MA) at the *NdeI* and *XhoI* restriction sites and expressed in *E. coli* BL21 (DE3). All the strains were maintained as frozen glycerol stocks.

### Seed conditions and media composition

The culture was revived from glycerol stock into 5 ml Super optimal broth [g/l: tryptone 20, yeast extract 5, NaCl 0.5, KCl 0.186], and was further transferred to 100 ml defined medium containing 5 g/l glycerol in 500 ml shake flask and incubated at 32°C and 180 rpm for 9 h. The defined media used for the seed and bioreactor cultivations were adapted from Korz et al. (1995) and contained glycerol as the sole source of carbon. The initial medium in the bioreactor contained [in 1l deionized water]: KH_2_PO_4_ 13g, (NH_4_)_2_HPO_4_ 4g, MgSO_4_.7H_2_O 1.2g, Citric acid 1.7g, EDTA 8.4mg, CoCl_2_.6H_2_O 2.5 mg, MnCl_2_.4H_2_O 15 mg, CuCl_2_.2H_2_O 1.5 mg, H_3_BO_3_ 3mg, Na_2_MoO_4_.2H_2_O 2.5mg, Zn(CH_3_COO)_2_.2H_2_O 13 mg, Fe(III) citrate 100 mg and thiamine 4.5 mg. Trace metals were filter-sterilized and added after autoclaving. Glycerol (5 g/l in seed, 15 g/l in production medium) was autoclaved separately and added before the start of the fermentation.

### Shake flask optimization

Preliminary studies were performed in shake flasks using the batch medium to establish the baseline growth characteristics and nitrilase activity. The medium was inoculated (5% v/v) with the seed culture and incubated at 32°C and 180 rpm. The concentration of β-D-isopropylthiogalactopyranoside (IPTG) and time of induction were also optimized.

### Fedbatch cultivation in bioreactor

Fed-batch cultivations were performed in a 3 l capacity Applikon bioreactor (Applikon Biotechnology, Delft, Netherlands) at 32°C, with agitation rate between 800-1200 rpm and aeration rate of 1 vvm (volume per volume per minute). The pH was maintained at 7.0 using 25% ammonia water and 1 M H_2_SO_4_. The bioreactor was filled with 800 ml initial medium, and inoculated (5% v/v) from the seed culture.

Dissolved oxygen (DO) concentration in the bioreactor was monitored using an autoclavable low drift AppliSens DO probe (Applikon Biotechnology, Delft, Netherlands). The overall oxygen mass transfer coefficient (k_L_a) for the bioreactor was determined using the dynamic gassing out-gassing in method (Nigam et al. 2012), by measuring the dissolved oxygen concentration (c) as a function of time in the equation ln (c^*^/c^*^-c) = k_L_a. t, where c* corresponds to saturation concentration. Minimum DO levels were set at 40% throughout the fermentation. DO control was achieved by cascading the agitation rate from 800 rpm to 1200 rpm followed by blending appropriate levels of pure oxygen in the air sparged (in the fed-batch phase). Foaming was controlled by with antifoam Y-30 emulsion (Sigma).

The initial batch phase for cell growth lasted 8-9 h; glycerol feeding was initiated after the DO level increased beyond 40% saturation after the initial drop to zero in the batch phase. The feeding solution contained 750 g/l glycerol, 15 g/l MgSO_4_ and trace elements and thiamine as in the initial medium. (NH_4_)_2_HPO_4_ (20% w/v) was applied as 10ml shots at appropriate intervals to keep the concentration of nitrogen and phosphorous at 50-200mM.

### Analytical methods

Samples were aseptically withdrawn from the culture after every 2 h from the start of the fermentation and used for the measurement of OD_600_ and analysis of dry cell weight (DCW, g/l). Amount of glycerol (g/l) and acetate (g/l) in the culture supernatant were estimated using biochemical estimation kits from Megazyme (Wicklow, Ireland), according to the manufacturer’s instructions. Ammonia and phosphate concentrations in the culture supernatants were measured using methods as described by Weatherburn (1967) and Chen et al. (1956), respectively. Biomass DCW was determined from washed cell pellets of 1ml culture dried overnight at 65°C. Intracellular protein production levels were monitored on 10% SDS-PAGE gel using standard methods (1970).

### Determination of enzyme activity

Nitrilase activity was determined in whole cells as described by Sohoni et al. (2015). Cells from 1ml culture were resuspended in 0.1 M sodium phosphate buffer (pH 7.5). Mandelonitrile stock solution (100 mM) was prepared in methanol. The reaction mixture (1 ml) contained 50 mM Tris–HCl buffer (pH 7.5), 10 mM mandelonitrile and 3-5 mg.DCW cells. After incubation at 37°C for 30 min, the reaction was stopped by the addition of 500 μl methanol. After centrifugation, the supernatant was analyzed for ammonia released during the reaction using phenol hypochlorite assay (Weatherburn 1967).

ADH and FDH activities were assessed in cell lysates by determining the rate of consumption of NADPH and NAD, respectively on a UV-visible spectrophotometer (Shimadzu, Kyoto, Japan). Absorbance was measured continuously in a quartz cuvette for 5 minutes at 340 nm. Acetophenone (10mM) or formate (20mM) was used as substrate for ADH and FDH, respectively.

### Results and Discussion

The production of industrially significant enzymes is routinely achieved through their expression in recombinant *E. coli* using high cell density cultivation. For instance, the cultivation of *E. coli* expressing nitrilases from *P. putida* (Naik et al. 2008), *Alcaligenes* sp. (Liu et al. 2011) and *P. fluorescens* (Sohoni et al. 2011) has been reported. However, studies on high cell density fed-batch cultivation for recombinant protein production in *E.coli* (including the above-mentioned reports) often use complex or semi-defined media, or generally include a yeast extract component in the feeding solution. The media composition and cost are key factors to be considered in fermentation (Shiloach and Fass 2005). Complex media provide easily digestible sources of nutrients and trace elements; it has also been shown that cheaper materials such as corn steep powder could be used (Liu et al. 2011). However, synthetic mineral media (besides low cost) are amenable to finer control of the specific growth rate and can deliver enhanced biomass yields with appropriate supplementation of nutrients (Shiloach and Fass 2005; Krause et al. 2011; Kangwa et al. 2015) In the present study we report a robust and simple to implement protocol based on fed-batch cultivation on a fully defined mineral medium, to achieve high cell densities of recombinant *E. coli* expressing the *Alcaligenes* sp. ECU0401 nitrilase with significantly high expression yields and enzyme activity. This nitrilase enzyme has been reported (Zhang et al. 2010) to be thermostable with excellent conversion of mandelonitrile to mandelic acid, making it highly attractive for industrial applications. The process described in the present study for HCD cultivation of *E. coli* and nitrilase production involves an initial batch phase with cell growth on glycerol in the medium, followed by a non-induced fed-batch phase with tightly regulated feed control, and finally a phase of sustained protein synthesis after induction with IPTG. Besides nitrilases, we also suggest that this method can be adapted as a general ‘plug and play’ protocol that can be applied for the large scale production of a range of enzymes in *E. coli* BL21/pET system without the need for separate optimization of the fermentation process.

### Shake flask optimization

Preliminary studies estimating activity on minimal medium and the optimization of induction conditions were performed on shake flasks. The concentration of IPTG to be used for induction was optimized at 0.2 mM (248.5 U/l). Higher concentrations (0.5 mM IPTG) led to reduction in growth rate and enzyme yield (136 U/l). Optimization of the proper conditions for induction is important for development of any method for recombinant protein production, especially in *E. coli* systems where strong promoters like T7 are used. The sudden onset of rapid protein synthesis has been suggested to impose a metabolic burden on the culture (Rosano and Ceccarelli 2014). In the current study, induction in the mid-log phase (OD = 1) after 3-4 h growth was observed to be optimum for sustained protein synthesis and maximum nitrilase productivity in shake flasks (248.5 U/l).

### Cultivation in bioreactor and modulation of Oxygen transfer rate (OTR)

Fig. 1 represents the features of a typical bioreactor run. When cultivated in a bioreactor, the culture grew to OD600~20 in the batch phase at a specific growth rate (μ) of 0.45 h^-1^. To satisfy the oxygen demand of the culture, the agitation speed was increased from 800 to 1200 rpm in the batch phase when the DO dropped below 40% (around 6h). Further increase in OTR was achieved by blending of pure oxygen at 100 ml/min initiated from the start of the fed-batch phase. Oxygen blending was increased in steps of 100 ml/min every hour till the induction of protein synthesis. In addition, it was observed that the overall oxygen mass transfer coefficient (k_L_a) of the bioreactor vessel (Fig. S1, Supplementary Information) varied more due to change in the agitation speed than the rate of aeration. Higher rates of aeration also led to escalation of foaming during the post-induction phase; therefore, the level of aeration was maintained at 1 vvm (volume per volume per minute) throughout the run.

**Fig. 1.**
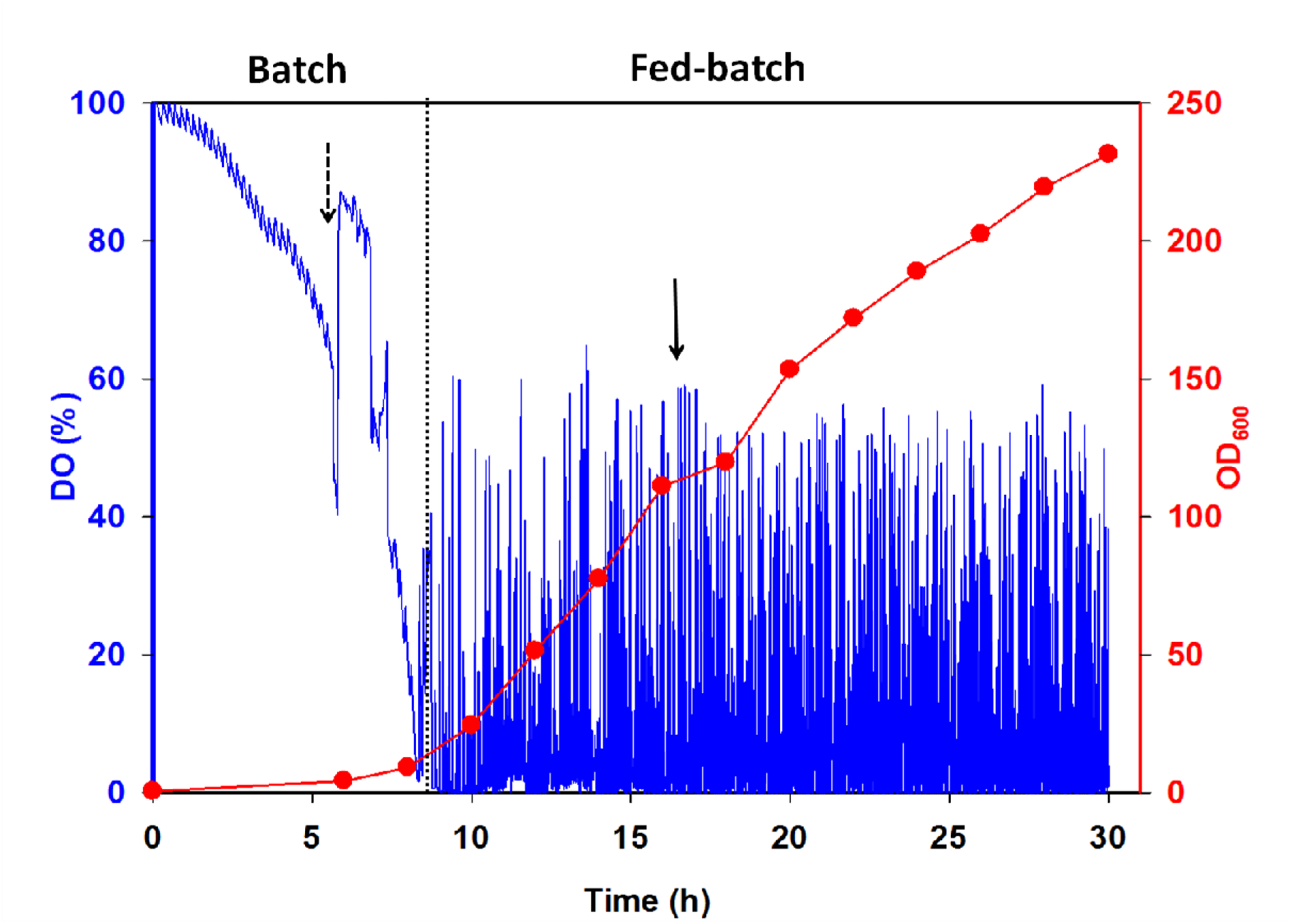
Production of nitrilase from HCD cultivation of *E. coli*: Example of the optimized fed-batch process applied in this study. Time points concerning increase in agitation speed (dashed arrow), start of feeding (dotted line) and IPTG induction (solid arrow) are indicated.

### Fed-batch strategy and enzyme production

The adoption of a properly controlled feeding approach is essential for the growth of *E. coli* to high cell densities and increased protein productivity (Shiloach and Fass 2005). It is desired to keep carbon as the growth-limiting nutrient, to avoid the formation of toxic by-products as a result of overflow metabolism. In the cultivation of nitrilase-expressing *E. coli*, feeding was started immediately after the glycerol and any organic acids formed in the batch phase were exhausted, as indicated by a sharp rise in DO from zero to above the 40% set-point. While trace elements and MgSO_4_ (15 g/l) were included along with the carbon source glycerol (750 g/l) in the feeding solution, (NH_4_)_2_HPO_4_ (200 g/l, 10 ml) was fed as concentrated shots at optimized time intervals to supplement nitrogen and phosphorous in the medium. Along with the ammonia fed for pH adjustment, this helped maintain the concentration of nitrogen and phosphate at 50-200mM, non-limiting yet non-toxic for cell growth, while preventing the precipitation of media components. Collins et al. (2014) have reported the use of separate carbon and N/P feeds for the HCD cultivation of recombinant *E. coli* expressing silk elastin-like protein.

Different carbon feeding strategies including constant feeding, exponential feeding and DO-stat have been used for production of enzymes through high cell density fed-batch cultivations. While constant or exponential feeding rely on preset feed or growth rates, DO-stat is a feedback control approach where feeding is controlled by the changes in dissolved oxygen concentration in the culture (Castan and Enfors 2000; Konstantinov et al. 1990). A small amount of glycerol from a concentrated feed solution is fed at each starvation signal, i.e. when the DO rises above the user-defined set-point due to depletion of the carbon source in the medium. The DO-stat method is therefore more responsive to the real time metabolic situation of the cell (Farell et al. 2015), and doesn’t involve iterative process development as in the case of exponential feeding. In defined media, DO-stat also responds faster to nutrient depletion than other indirect feedback control strategies such as pH-stat (Lee et al. 1996, Picotto et al. 2017). Based on these advantages, the DO-stat feeding approach was chosen for this study.

During the fed-batch phase of the fermentation, the DO values were observed to oscillate between 0-60% (Fig. 1) around the 40% set-point. DO values greater than the set-point indicated momentary starvation phase, while lower values suggested momentary overfeeding. The overshooting possibly occurs due to limitations of the hardware employed, such as a probable time delay of signal from the DO probe (Lee et al. 1994), and the fact that feed flow rates of as low as 10-50 ml.h^-1^ need to be maintained for working reactor volume of ~ 1 l.

With the DO-stat method, we further observed that the modulation of PID (proportional, integral and derivative) controller settings integrated in the bioreactor also helped fine tune the feeding after every starvation signal. Often, due to the complexity and variability of microbial fermentations, the optimum PID settings are required to be determined through empirical methods (Potvin et al. 2012). The default PID settings (7:1500:1) were aggressive (larger proportional gain), and led to increased amount of glycerol feeding per signal, shorter starvation periods and a faster growth rate. On the other hand, reducing the proportional grain to a moderate level (3:1500:1) helped feed less glycerol after every starvation period, as demonstrated by the closer spacing of the DO oscillations (Fig. 2). With the moderate PID settings, 0.3-0.4 g/l glycerol was fed after each starvation signal, which was immediately used by the growing cells. This strategy afforded the maintenance of low levels of glycerol in the medium during the fed-batch phase, thereby also preventing the accumulation of acetate in the medium (Fig. 3a). Further, in an oxygen limited culture, the specific growth rate (and in turn, the rate of uptake) is proportional to OTR. The glycerol uptake matches the glycerol feed rate, as no glycerol ideally accumulates in the medium in the DO-stat mode. Thus, glycerol-feeding rate (F_glycerol_), and by extension the rate of cell growth, could be readily controlled by varying the OTR in the DO-stat mode of operation. The feed rate showed a near-linear dependence on OTR, suggesting the absence of overflow metabolism (Fig. 3b). Relatively high μ values of 0.25-0.36 h^-1^ were maintained in the early part of fed-batch phase, up to cell densities of 35 g DCW.l^-1^. In fact, we could achieve a biomass concentration of 98 g DCW.l^-1^ in 24 h when the culture was grown at 32° C without induction (data not shown).

**Fig. 2.**
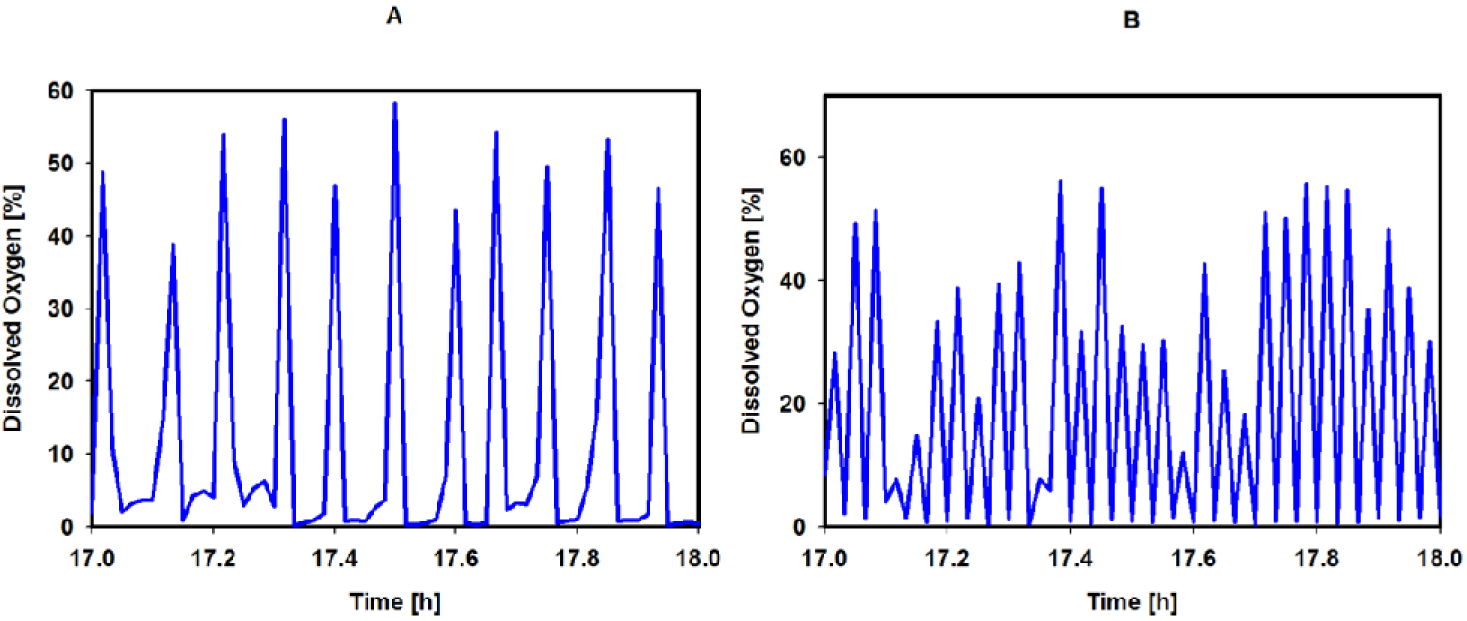
Effect of (a) aggressive and (b) moderate PID controller settings on the dissolved oxygen (DO) profile in DO-stat based fed-batch strategy

**Fig. 3.**
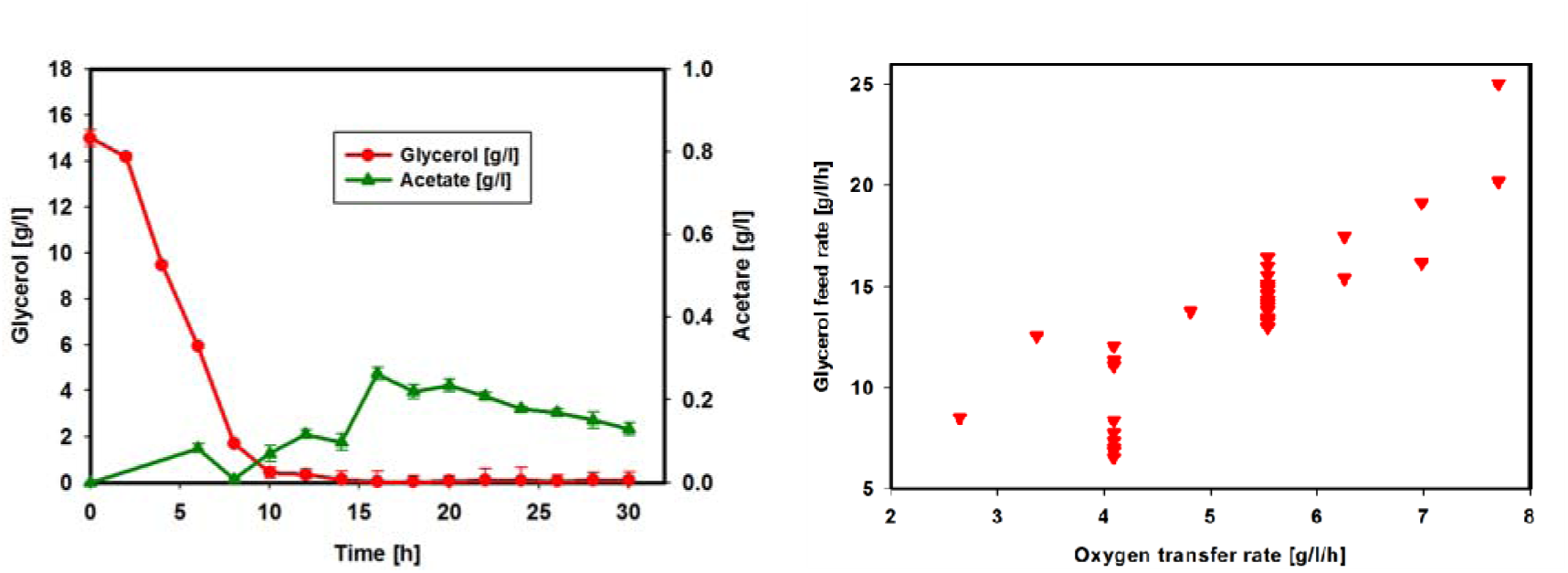
(a) Time course measurement of glycerol and acetate concentrations in a representative fed-batch run (b) Dependence of growth and glycerol uptake rates on oxygen transfer rate (OTR) in the bioreactor vessel

Protein production was induced at the mid-log phase OD~100 with 0.2mM IPTG, as determined in shake flask experiments for better nitrilase productivity. It should be noted that the use of fully defined media ensures the absence of any leaky expression, allowing the cells to grow to moderately high densities with minimal metabolic burden. We also tested the addition of yeast extract as complex additive at induction (“boosting”), which has been suggested to enhance protein synthesis and folding (Krause et al. 2010; Tsai et al. 1997). However, the introduction of 2% (w/v) yeast extract did not enhance the enzyme production; rather, it stalled the growth of the culture and led to excessive foaming (data not shown).

The choice of feeding strategy also plays a critical role during the protein synthesis phase. In the DO-stat approach, the culture is constantly starved of carbon substrate throughout the fed-batch phase, thereby preventing overfeeding and directing more energy towards recombinant protein production. We observed that accidental overfeeding of glycerol in our reactor runs usually led to decreased enzyme productivity in spite of normal growth, with the repression of protein production extending even after the overfed glycerol was used up (Fig. S2, Supplementary Information).

Induction of protein synthesis at mid-log phase (OD_600_ ~ 100) along with the DO-stat feeding approach yielded up to 140 kU/l nitrilase enzyme titer (Fig. 4), which already represents a 7-fold increase over the maximum productivity of *Alcaligenes* sp. nitrilase (19.3 kU/l) reported earlier (Liu et al. 2011). However, the culture did not sustain at high growth rates beyond 24 h at 32°C; excessive foaming and cell lysis (denoted by drop in OD) were observed. In order to further increase enzyme yield, we employed a reduction in temperature, coupled with the lowering of OTR, to achieve an optimal growth rate conducive for sustained protein synthesis and proper folding.

**Fig. 4.**
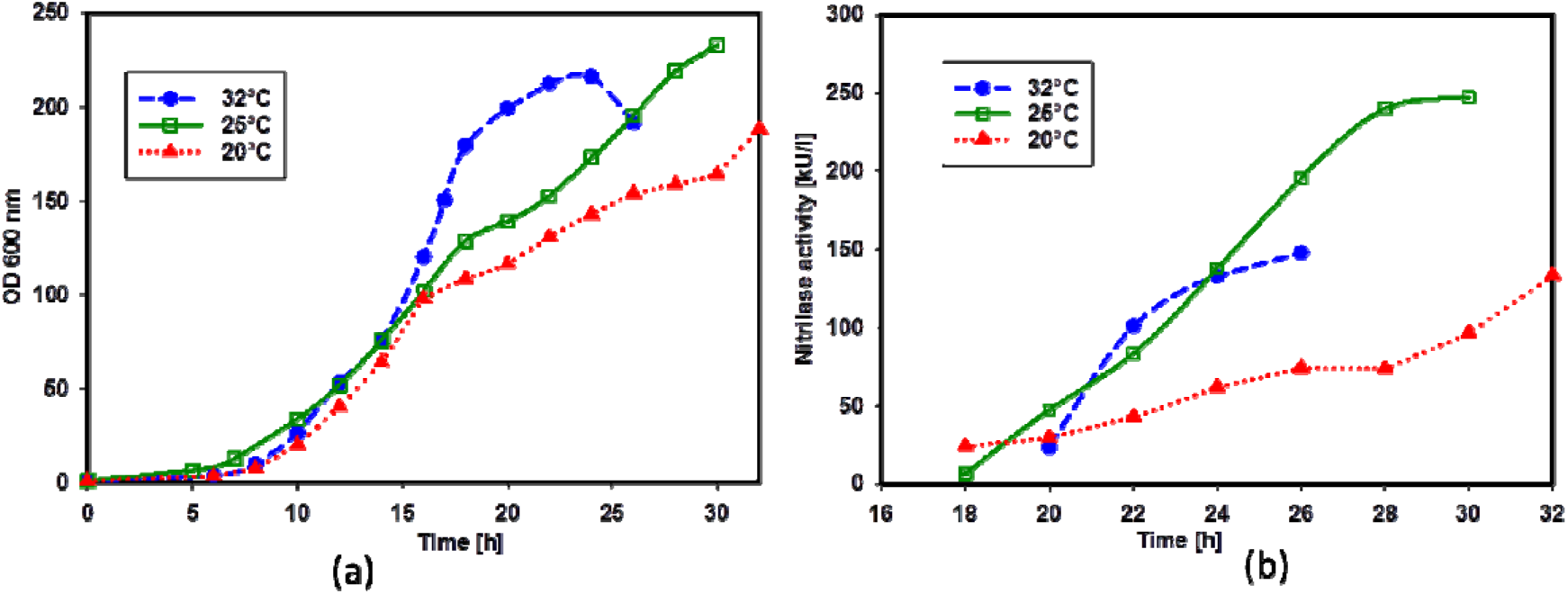
Effect of post-induction temperature on growth and enzyme titer. *E. coli* BL21 (DE3) expressing *Alcaligenes* sp. nitrilase was grown at 32°C in a fed-batch mode till around 16 h, when the cell density reaches OD_600_~100 (35 g DCW/l). The culture was then induced with 0.2 mM IPTG and the temperature was either kept unchanged at 32°C or reduced to 25°C or 20°C. (a) OD_600_ and (b) nitrilase activity (kU/l) profiles for the corresponding reactor runs

The optimized protocol involved growth up to 35 g DCW.l^−1^ at 32°C by maintaining a relatively high growth rate of 0.23-0.36 h^−1^ in the fed-batch phase (Fig. 5), with peak OTR at ~7.5 g O_2_.l^−1^.h^−1^ just prior to induction. At the time of induction, the temperature was reduced to 25° C with concomitant lowering of OTR to about 5.5 g O_2_.l^−1^.h^−1^ (500 ml/min oxygen blending) and continued till 24 h, after which the OTR was further lowered to ~ 4 g O_2_.l^−1^.h^−1^ (300 ml/min oxygen blending). The growth rate dropped substantially to under 0.1 h^−1^ as a result of this process manipulation. Using this strategy, protein synthesis could be sustained for a longer period (10-12 h), resulting in a 1.7-fold increase in enzyme yield to 247 kU/l (Fig. 4). A trade-off was observed between growth rate and soluble enzyme production; a higher growth rate and cell mass ensued at 32°C but higher enzyme titres were achieved at 25°C. Liu et al. (2011) have also observed that higher cell mass do not necessarily relate to increased nitrilase titers. However, a further reduction in temperature to 20°C appeared to slow down the rate of growth and protein synthesis, resulting in much lower enzyme yields (Fig. 4).

**Fig. 5.**
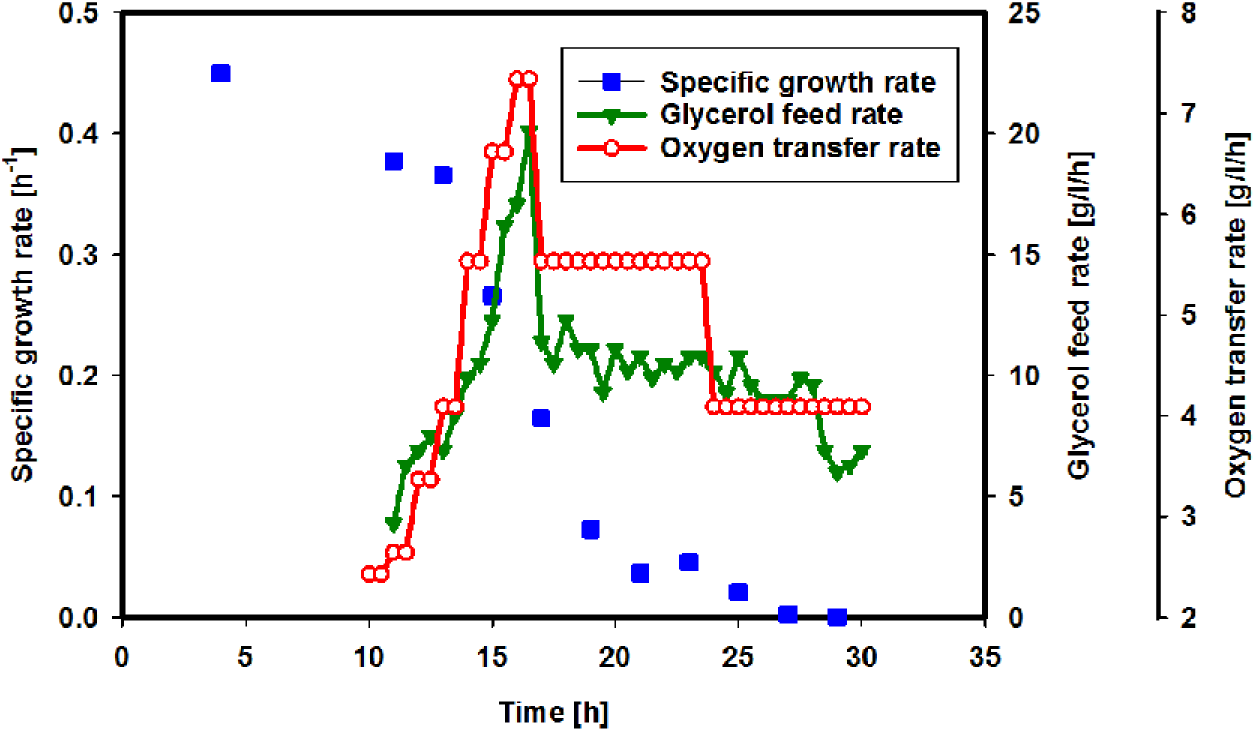
Time course measurement of specific growth rate (h^-1^), glycerol uptake rate (g.l^-1^.h^-1^) and OTR (g.l^-1^.h^-1^) for a representative (optimized) fed-batch process. The OTR was varied by varying the speed of agitation and oxygen blending

### Final biomass and enzyme yields

The final optimized process yielded OD600 = 234 (cell mass 78 g DCW/l) after 30 h with an enzyme titer of 247 kU/l (Fig. 4). Based on the elemental composition of *E. coli* biomass from literature (Roels 1980), the overall biomass yield coefficient achieved using the fed-batch process was calculated to be 0.36 g DCW/ g glycerol fed in the medium, or 0.44 C-mol biomass/ C-mol glycerol. The soluble expression and enhancement of enzyme production was also confirmed on SDS-PAGE (Fig. S3, Supplementary Information). From the known specific activity of purified *Alcaligenes* sp. nitrilase (25.6 U/mg, Zhang et al. 2010), this enzyme yield corresponds to ~9.6 g/l soluble protein. To our knowledge, these values are the maximum reported for any nitrilase enzyme, and manifold greater than the previous titer and cell density achieved (Liu et al. 2011) for the recombinant *Alcaligenes* sp. nitrilase (19.3 kU/l with 10.7 g/l DCW, through constant feeding of glycerol at 9 g/h in complex media). In addition, it has been reported (Zhang et al. 2010, Liu et al. 2011) that the soluble overexpression of the recombinant *Alcaligenes* sp. ECU0401 nitrilase in *E. coli* JM109/pUC19 is superior over the BL21/pET11a or pET28a system. However, using the optimized fed-batch strategy reported here, we were able achieve enhanced high cell density and titers of soluble active enzyme even with the BL21/pET expression system. We believe that the key to achieving high enzyme titers resides in the tight control of the bioprocess, through the use of fully defined media, application of the DO-stat feeding strategy and modulation of post-induction growth rates for sustained protein synthesis. The robustness of the process can be improved further with the implementation of better feedback controllers and automated reactor systems.

### Applicability for other enzymes

The applicability of the strategy described in this study was tested for the HCD cultivation of recombinant *E. coli* BL21 (DE3) expressing two other well-characterized enzymes of commercial relevance, alcohol dehydrogenase (ADH) and formate dehydrogenase (FDH). ADH from *L. kefir* (Hummel 1990) has been used in the industry for bioreduction of carbonyl to hydroxy group in a variety of ketone substrates, including acetophenone to (R)-phenylethanol. Using the optimized bioprocess from the present study, an ADH productivity of 7115 kU/l was achieved with cell growth up to OD_600_ = 248, which corresponds to 12.9 g/l protein. Meanwhile, the *C. boidinii* FDH finds application as an effective NAD cofactor recycling enzyme (Tishkov and Popov 2006) in biotransformations involving oxidoreductases (including ADH). The *C. boidinii* FDH has been used on an industrial scale (Degussa AG) for co-factor regeneration in the production of L-*tert-* leucine from trimethyl pyruvate catalyzed by leucine dehydrogenase (Breuer et al. 2004). The optimized fed-batch process applied for FDH production gave an enzyme yield of 48 kU/l (Fig. S4, Supplementary Information) with cell growth up to OD_600_ = 275.6. The improvement of enzyme yields due to reduction in temperature and OTR was more significant in the case of FDH (48 kU/l at 25°C vs 6.6 kU/l at 32°C). It is significant that we could achieve excellent yields for these enzymes using the DOstat fed-batch process developed in this study with minimal modifications, which proves the effectiveness of the method for application in industrial scale production of active recombinant enzymes.

## Conclusion

In conclusion, the fed-batch strategy demonstrated here presents a simple scalable method for recombinant protein production in *E. coli* high cell density cultivations on defined media. With strict control on the growth rate achieved via DOstat nutrient feeding and appropriate process manipulations, maximum cell growth and enzyme yields have been demonstrated in the case of nitrilase, ADH and FDH. This method further provides a reliable template which can be used on a benchtop fermentor to produce large amounts of soluble and active enzyme that could be readily used for biotransformation trials en route developing commercial biocatalytic applications. Although the method was developed with known industrially significant enzymes that express well in *E. coli*, we suggest that it could also perform well for other related applications such as the production of high-value pharmaceutical proteins and the cultivation of metabolically engineered *E. coli* strains for the production of platform chemicals.

## Declarations

### Authors’ contributions

PPW and RPG conceived the research. PPW, SVS and RPG designed the experiments. SVS, PHK and AGS performed the experiments and analyzed the data. AVS, SVS and PPW wrote the manuscript. All authors read and approved the final manuscript.

## Acknowledgement

Technical assistance provided by Shikha Shah, Annesha Sengupta and Swati Madhu in performing the reactor runs and biochemical assays is duly acknowledged.

## Consent for publication

All authors have read and approved to submit it to Biorxiv. There is no conflict of interest of any author in relation to the submission.

